# Semantic map learning and externalization in an embodied neural agent: A comparison to human behavioral and neural data

**DOI:** 10.64898/2026.05.19.726370

**Authors:** Kathryn Simone, Lea Steffen, Nicole Sandra-Yaffa Dumont, Siyang Yu, Hudson Ly, Graeme Damberger, Chris Eliasmith

## Abstract

All mammals can build internal cognitive maps from sensory input, supporting spatial learning and planning. While rodent studies have shown the mammalian navigation system solves the SLAM (Simultaneous Localization and Mapping) problem, its role in human spatial memory and overt recall are less understood. To investigate this, we adapted a spiking semantic SLAM algorithm for a “Treasure Hunt” task, where human participants navigate a 3D beach in virtual reality and later point to remembered object locations. Our agent integrates networks for bipedal locomotion, vision, memory, and arm control to enable first-person learning of place-object associations, and externalizing that knowledge by pointing and expressing confidence. Comparing model observables to human data, we replicate key behavioral and neural effects: monotonic scaling of accuracy with confidence, and recall-dependence on local field potential power observed in the left hippocampus. This work offers a mechanistic framework linking embodied navigation, memory, and communication in human spatial cognition.

## Introduction

Both humans and animals rely on knowledge of their environments to navigate, forage, and avoid danger within their ecological niches. In mammals, spatial cognition is distinctly prominent: by integrating raw sensory experience, animals can synthesize an internal allocentric map — a bird’s-eye view — to represent the spatial relationships among salient environmental features. This concept of a *cognitive map* was proposed by Tolman (1948) to explain rodents’ capacity for flexible navigation, such as planning efficient shortcuts to goal locations. This sophisticated mapping stands in contrast to insect navigation strategies that can often be captured by vector-based homing or route-following heuristics (e.g., bees and ants; Dyer, 1991; Wehner et al., 1990, 2006). Human spatial cognition includes the capacity to *externalize* internal spatial representations through gesture and other modalities. This allows spatial knowledge to support collaborative goals, such as cooperative navigation and sharing of knowledge about unseen locations. This is critical to the performance of the collective, as evidenced by evolutionarily-convergent mechanisms for gesture-like communication of spatial knowledge in other species, including bee waggle dances (Seeley & Buhrman, 1999).

Across mammalian species, navigation relies on shared substrates in the hippocampal-entorhinal system, including those that exhibit grid-like coding and path integration observed in rodents (Benhamou, 1997; Hafting et al., 2005) and bats (Aharon et al., 2017; Yartsev et al., 2011). Analogous computations have also been identified in humans (Doeller et al., 2010; Jacobs et al., 2013; Mittelstaedt & Mittelstaedt, 2001; Zhu et al., 2023). A functional interpretation of these processes is possible through the computational formalism of *simultaneous localization and mapping* (SLAM): the computational process of concurrently estimating one’s position (localization) and constructing a spatial representation (mapping; Durrant-Whyte and Bailey, 2006) while moving through an unfamiliar environment. This is a well-understood problem in robotics, where the need to build an internal map from first-person experience in a novel environment appears in home robotics, automotive applications, and commercial navigation (Taheri & Xia, 2021).

Facing similar demands, biological systems appear to have converged on analogous solutions. The rodent hippocampal-entorhinal system, for instance, has been interpreted as a biological SLAM-like system that integrates self-motion (path integration) with landmark cues to simultaneously localize and map environments (Safron et al., 2022). A suite of biologically-inspired SLAM algorithms emphasize the repurposing of this general neural solution in service of the particular embodiment, including RatSLAM for robust 2D navigation (Milford et al., 2004), BatSLAM and DolphinSLAM for sonar-based 3D localization (Silveira et al., 2015; Steckel & Peremans, 2013), and NeoSLAM for long-term visual SLAM using structured spatiotemporal representations inspired by human memory (Pizzino et al., 2024). By extension, while humans inherit the neural circuits underlying spatial cognition that evolved in mammals, their role in the externalization behaviors that distinguish us from other animals is unclear. Multi-agent SLAM systems demonstrate that spatial collaboration—similar to that observed in humans—requires representational forms that are both internal *and* communicable (Lajoie et al., 2022), a challenge not addressed by existing models of the hippocampal-entorhinal system. Recent robotics architectures such as LM-Nav demonstrate how an agent’s internal spatial knowledge can be externalized into language, enabling users to specify goals via natural-language landmark descriptions that are directly grounded in agent perception (Shah et al., 2023).

Despite their functional power, these approaches are not designed to yield biologically grounded, experimentally falsifiable hypotheses about the neural mechanisms by which humans transform, appraise, and communicate spatial knowledge. Consequently, there remains no unified computational account of how conserved mammalian navigation systems are repurposed to meet the symbolic, probabilistic, and socially communicative demands unique to human spatial cognition.

Neurally implemented *Vector Symbolic Algebras* (VSAs) offer a promising framework to unify spatial memory, subjective appraisal, and motoric externalization—in a manner grounded in neural structure suitable for experimental modeling. VSAs bridge symbolic and connectionist approaches by projecting structured representations into a hyperdimensional vector space where algebraic operations support the composition and manipulation of information, in a manner that can be mapped to neural networks. An architecture that does so for spiking neural networks is known as the Semantic Pointer Architecture (SPA; Eliasmith, 2013). The preferred VSA in the SPA is Holographic Reduced Representations (HRRs; Plate, 1992). While other algebras have been proposed, the HRR algebra admits representing continuous structure via *fractional binding*, a property important for map encoding and leveraged by *Spatial Semantic Pointers* (SSPs; Komer et al., 2019).

SSPs jointly encode semantic content and allocentric spatial structure, enabling unified representations of objects, locations, and relations that can be queried, transformed, compared, and communicated. These operations align naturally with the computational demands of human spatial reasoning in social contexts. Importantly, because the SPA has shown how to map HRRs to spiking networks, and mammalian neuroanatomy more generally, VSA data structures admit direct biological interpretations. For example, it has been demonstrated in Dumont and Eliasmith (2020) that continuous spatial states can be encoded as hexagonal SSPs (HexSSPs), spiking responses directly map onto the grid-cell-like firing patterns observed in the entorhinal cortex.

Building on this, Dumont et al. (2023) introduced SSP-SLAM, a fully spiking SLAM system that leverages grid-cell-based SSP representations to perform localization and mapping under biologically realistic constraints, providing a concrete mechanistic account of how hippocampal–entorhinal circuitry could implement semantic SLAM-like computations while building the representations needed to manipulate spatial knowledge.

Beyond navigation, VSAs provide a route to modeling the *probabilistic appraisal* of internal representations. Furlong and Eliasmith (2022) established formal correspondences between VSA similarity operations and probabilistic inference, grounding vector similarity in probability statements. This provides a natural interpretation of subjective confidence as a similarity-derived quantity computed over distributed neural representations. This connection suggests a unified account in which the same representational substrate supporting spatial memory and localization also enables confidence estimation and communication—capacities essential for the social use of spatial knowledge in humans.

Together, these developments motivate the hypothesis that VSAs—implemented through SSPs and grounded in spiking grid-cell population codes—provide a unifying computational substrate through which evolutionarily ancient navigation mechanisms are repurposed to support the probabilistic evaluation, externalization, and social communication distinctive to human cognition.

In this paper, we integrate SSP-SLAM into an embodied cognitive model of human spatial learning, offering a mechanistic account of how an evolutionarily ancestral navigation algorithm is adapted for human social contexts. We focus on the Treasure Hunt task (Miller et al., 2018), which requires participants to encode, remember, and communicate spatial information, and for which both behavioral measures and direct neural recordings are available. Importantly, this task motivates an embodied cognitive model because it requires: *(1)* integration of self-motion cues with visual landmark perception during active navigation, the core computational challenge addressed by SLAM; *(2)* formation of a stable learned spatial map that is accessible after exploration ends; and *(3)* transformation of internal spatial representations into externally communicable responses. We evaluate the model by testing whether it reproduces two key human signatures from this task: behavioral confidence–accuracy scaling and a memory-related effect low-theta activity in the left hippocampus.

## Mathematical Preliminaries

### Spatial Semantic Pointers (SSPs)

SSPs represent continuous variables, such as those that define spatial location *x* ⊆ ℝ^*m*^, in a *d*-dimensional space *X*⊆ ℝ^*d*^ via a Fourier-domain construction,

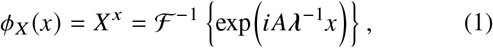

where *λ* is a non-negative diagonal matrix setting length-scales, and *A* ∈ ℝ^*d*×*m*^ is a random *phase matrix* that is related to the ‘axis vector’ X via the inverse Fourier transform. For mapping a 2D space, *m* =2. The columns of *A* are built from collections of frequencies and are constrained to have conjugate symmetry so that *ϕ*_X_(·) is real-valued. This construction ensures unit-magnitude Fourier components (i.e., unitary vectors), which is important for stable iterative composition. Symbol-like discrete data, such as that which encodes object identities, are also represented as unitary vectors. In this work, we adopt the common strategy of assigning to each such symbol-like discrete value a random unitary vector such that distinct symbols are approximately orthogonal, i.e., *ϕ*(*i*)·*ϕ* (*j*) ≈0 for *i* ≠ *j*. We refer to symbol-like discrete representations as Semantic Pointers (SPs), and follow the convention of Komer et al., 2019 to refer to the continuous representations as Spatial Semantic Pointers (SSPs).

Notably, both SSPs and discrete representations use the HRR operations implemented in spiking neurons. Specifically, the *binding* operation (circular convolution, ⊛) is used in two ways: 1) to generate a composite vector from two symbol-like inputs; and 2) as a recursive operation to define a point on a continuous axis, e.g. *ϕ*_*X*_ (*x*) = *X*^*x*^. For spatial cognition, binding is applied to associate the SP representation of an object, *B*, with the SSP-represented location in the environment, 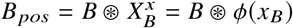.

Accessing one item of the pair of vectors that form the bound product is achieved through *unbinding*, which is implemented by binding the composite representation with the involution of the other item in the pair, e.g., (*B* _*pos*_ ⊛ *B*^−1^ ≈*ϕ* (*x*_*B*_), where ^−1^ denotes the involution operation. Unbinding allows querying of encoded associations.

Importantly, due to the properties of circular convolution of unitary vectors, binding and unbinding of SSPs maintain a correspondence to the original metric space (Komer, 2020). Specifically, binding of two SSPs produces a new SSP corresponding to the representation of the sum of the encoded variables: *ϕ* (*x*_1_) ⊛ *ϕ* (*x*_2_) = *ϕ* (*x*_1_ +*x*_2_). Similarly, unbinding reflects the difference: *ϕ* (*x*_1_)⊛ *ϕ*^−1^ (*x*_2_ = *ϕ* (*x*_1_−*x*_2_).

The *bundling* operator is used to group vectors into a single representation, and is superposition of representations via vector addition. While this operation is dimensionality-preserving, the output is generally not constrained to the set of valid SSPs, unitary, or even unit-norm vectors.

The final operator defining our vector algebra is *similarity* for which we use the inner product. Notably, the inner product of two SSPs defines an approximate sinc function of the distance between the encoded variables (Voelker, 2020): *k* (*x, x*^′^) = *ϕ*(*x*) *· ϕ*(*x*^′^) ≈ sin(*π*|*x* − *x*^′^ |) /(*π*|*x* − *x*^′^|). This kernel means that an SSP bundle 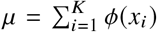can be interpreted as representing an empirical probability distribution over its constituent vectors (Furlong & Eliasmith, 2022). As a result, computing the probability of a new sample point, *x*, is equivalent to taking the inner product between the SSP probability density representation and the SSP encoding of that point *x*, through some function *f* : *p*(*x*) ≈ *f* {*ϕ* (*x*) · *µ*}. One suitable form for *f* is rectification with an appropriate bias (Glad et al., 2003). That is:

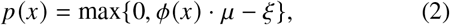

where *ξ* is fit on a per-distribution basis.

Given the compressive nature of bundling, binding, and unbinding operators, a suitable *cleanup* operation is sometimes used to recover a vector within the valid set. In our model we use *X* to perform cleanup (Komer, 2020).

### SSP SLAM

SSP-SLAM is a biologically-plausible spiking neural network implementation of Simultaneous Localization and Mapping (SLAM; Dumont et al., 2023). It uses SPA methods and exploits semantic information by binding SPs to SSPs to learn a cognitive map of the environment.

The network maintains a continuous estimate of the agent’s self-position, represented as an SSP 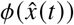, through path integration of velocity cues. The path integration network implements a hybrid oscillatory-interference and continuous attractor dynamics model, wherein position is tracked by ⌊*d* /2 ⌋ velocity-controlled oscillators (VCOs), one for each Fourier coefficient of the SSP. Each VCO implements self-stabilizing dynamics that maintain the unit circle as an attractor, preventing drift from the valid SSP manifold.

The map itself is learned with an associative memory. The associative memory uses spiking neurons with learned encoders and decoders to acquire place-object associations that define the cognitive map. These associations are learned online through the biologically plausible Voja (Voelker et al., 2014) and Prescribed Error Sensitivity (PES; (Bekolay et al., 2013)) learning rules. The Voja rule shifts incoming neuronal encoders toward frequently encountered inputs, while the PES rule modifies outgoing synaptic weights (decoders) in response to error signals.

Exploiting the HRR operations detailed above, the model performs key spatial transformations: binding with the self-position SSP converts egocentric landmark observations into allocentric map entries (*ϕ* (*x*_*ego*_) ⊛ *ϕ* (*x*_*sel*_) _*f*_ = *ϕ* (*x*_*allo*_)); and unbinding egocentric feature vectors from stored allocentric locations computes a second self-position estimate for loop closure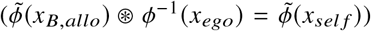. The cognitive map can be characterized as a bundle, effectively superimposing multiple object-location associations in a single, queryable representation. Critically, the use of spiking neural dynamics permits direct comparison with neural data recorded from biological systems — an analysis not feasible with non-spiking SLAM approaches.

### Embodied Cognitive Model

Embedding SSP-SLAM in a more complete agent model, our model demonstrates how an embodied agent can acquire and later express spatial knowledge in the context of the Treasure Hunt task (see Figure 1). Developed by Miller et al. (2018) this is a virtual reality experiment requiring participants to navigate a virtual beach environment and visit four randomly-positioned treasure chests, which open to reveal either an empty chest or a unique item. Following exploration, participants fly from their current position to an elevated vantage point. After completing a distractor game, participants are then queried about each item encountered, first to provide a subjective confidence report in whether or not they recall of the object’s location (Yes, No, Maybe), and then to position a target circle on the virtual beach. Performance is quantified using a rank-based error metric.

**Figure 1:**
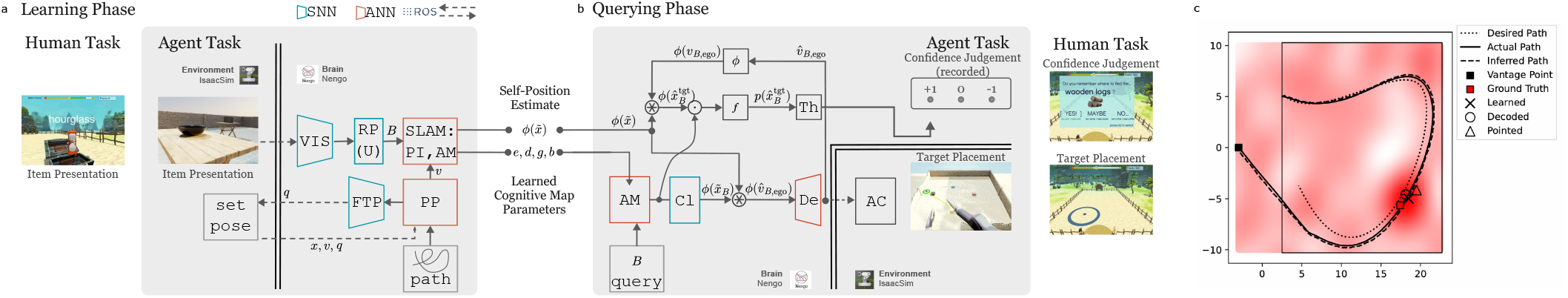
Architecture of the embodied cognitive model in the **a** Learning Phase and **b** Querying phase. Acronyms defined in the main text. **c** Depiction of model observables, showing correspondence between actual and reported landmark locations. Screenshots from human VR task adapted from Miller et al. Fig. 1, under CC BY 4.0. Changes: Cropped.

We addressed the demands of the Treasure Hunt task associated with the learning and querying phases separately, through two simulations linked through the SSP-SLAM network state that defines the agent’s cognitive map. The learning simulation (Fig. 1a) integrates vision (VIS), a random unitary projection layer (RP(U)), spatial memory (SLAM), and motor control (PP, FTP) to guide an H1 humanoid robot through a 3D simulated environment in IsaacSim, building a cognitive map in a spiking neural network. This system models how humans form spatial representations through experience. The querying simulation (Fig. 1b) uses the associative memory module (AM), that encodes the learned cognitive map with cleanup (Cl), decoder (De), scalar thresholding (Th), and arm control (AC) to produce two key behavioral readouts of the Treasure Hunt task: confidence judgments and pointing. Querying by-passes visual recognition, directly presenting symbolic place-object queries (query) to the associative memory module, and computes time-averaged confidence and motor outputs over a 3-second window. Together, the two simulations demonstrate a unified characterization of map learning and recall in embodied agents (Fig. 1c).

### Learning Phase

#### Path Generation and Following

For navigation, a waypoint generation algorithm computes paths between table/chest locations by spatially distributing targets and reordering them to minimize traversal distance through circular optimization around a geometric centroid. These discrete waypoints are interpolated using cubic spline smoothing to generate continuous reference trajectories (Fig. 1c). A bipedal walking robot follows the specified path through a neural hierarchical control architecture. A spiking implementation of a pure pursuit controller computes the error between the desired point along the path at timestep *t, x*_ref_ (*t*) and actual robot position *x* (*t*) to generate a desired forward speed and angular rotation. This is then provided as input to a non-spiking reinforcement-learning trained policy network optimized for flat terrain, which together with other state information is used to compute and set 19 joint angles *q* (*t*) that dynamically sets the robot’s pose to implement walking.

#### Vision

The vision system identifies objects for the H1 agent in IsaacSim using a YOLOv5s (Jocher, 2020) detector fine-tuned for 100 epochs from pretrained weights on simulated first-person RGB camera images. Classifications other than those of the top eight most reliably recognized objects were ignored by the network. Class labels corresponding to a recognized object among the selected were mapped to a semantic pointer *B*. When no recognized object was present, learning was suppressed.

#### Simultaneous Localization and Mapping

The spatial memory system implements SSP-SLAM as introduced in the SSP SLAM Section. The path integrator is initialized to the origin and updated by robot velocity *v*. We have integrated a spiking rectified flow cleanup network (Habashy, 2026) to maintain a valid SSP representation and track position robustly over time.

### Querying Phase

We formalize the querying phase problem as follows: Given item query *B*, self-position estimate 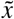, and cognitive map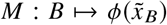, compute an egocentric vector *v*_*B*,ego_ ∈ ℝ^2^ such that 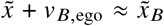, and a confidence judgment *c* ∈ {−1, 0, 1 }.

#### Pointing Vector

Presentation of object query *B* to the Associative Memory (AM) yields a recall embedding 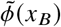, which encodes the remembered position of the object. This and the self-position SSP 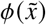 are passed through cleanup to produce valid SSPs 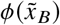and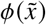. The egocentric displacement SSP *ϕ v*_*B*,ego_ is computed by unbinding: 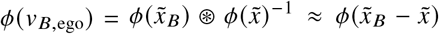. This SSP is decoded by a spiking ensemble approximating *ϕ*(*v*) ↦*v* over *v*∈ 𝒱, thedisplacement bounds defined by the environment.

#### Confidence Judgment

To obtain a confidence judgment from the model, we interpret the question “Do you remember where to find *X*?” as a request for the likelihood that the participant’s *next* pointing response will be “good”. We define a good response as placing the cursor within a reward radius *r* = 3 of the true allocentric landmark location, motivated by the sufficient condition for which subjects were awarded points in the human study (reward “disk” of radius 3 virtual units).

We hypothesize that human confidence reports reflect the probability density assigned to the *intended* allocentric target location under the AM-induced belief. In the model, recall induces a spatial belief over allocentric locations *x*∈ 𝒳 ⊂ ℝ^2^. We form an allocentric target estimate via binding: 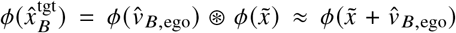 and compute the probability as per equation 2. Specifically,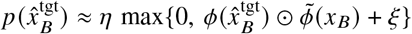 where *ξ* is the bias, selected so that the kernel is active (non-negative) for a proportion *ρ* = 0.6 of a representative set of similarity evaluations, and *η* is the normalizing constant computed at each timestep to ensure that 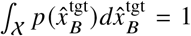. Finally, we map the probability density to the three-level confidence report *c* ∈ {−1, 0, 1} using two reference densities: the uniform baseline 1/*A* (where *A* = area(V)), and the characteristic density of a top-hat distribution supported on the reward disk, 1/(*πr*^2^):

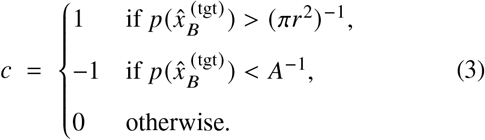

#### Arm Control and Cursor

The intended pointing vector *v*_*B,ego*_ is transformed to a target location within the Simulator’s coordinate system through summation with the robot’s current 2D position. The result defines a point on the *Z* = 0 plane. As control becomes unstable for targets beyond the robot’s reach, this target point is projected onto a reaching sphere centered at the robot’s shoulder. Pointing to this target is achieved through control of the four joints of the right arm: the shoulder pitch, shoulder roll, shoulder yaw, and elbow joints. The cursor location, which we take as the model output, was computed as the point on the floor that intersects with line between the elbow and hand joints. Arm control is implemented using the ABR Control package (DeWolf et al., 2020), adapted for the H1 humanoid in IsaacSim.

## Experiments and Results

Having shown that our model can effectively perform the Treasure Hunt task, we now assess how well its internal mechanisms replicate human data. By conducting side-by-side comparisons of three behavioral analyses and one neural analysis from Miller et al. (2018), we test whether our model can account for the spatial memory and hippocampal dynamics observed in humans. The original study emphasized characterizations of pointing accuracy and confidence. These analyses are reproduced in the top row of Fig. 2, and (a) reveal high intersubject variability in pointing accuracy, (b) suggest a correlation between confidence and accuracy on an individual basis, and finally (c) indicate similar confidence-conditioned probability density functions over accuracy between individuals. We adopt the definition for accuracy in the original paper,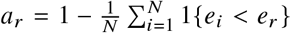 which computes the percentile rank of the error of the actual response, *e*_*r*_, among the set of *N* possible responses indexed over *i*, due to discretization.

**Figure 2:**
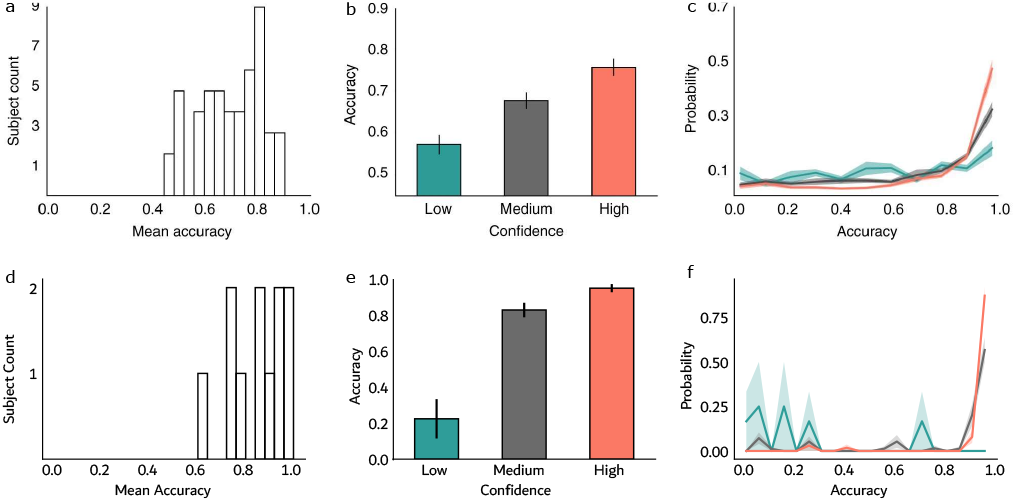
Behavioral data from human subjects (top row) and model (bottom row). **a** Histogram of mean accuracy across subjects (*N* = 46 subjects, *M* = 11 models). **b** Mean accuracy conditioned on confidence level. Error bars are ±SEM. **c** Accuracy distributions for each confidence level. Solid lines show mean density across individuals; shaded regions are ± SEM. Human data (top row) adapted from Miller et al. Fig. 1, under CC BY 4.0. Changes: Cropped.

To simulate intersubject variability, we instantiate a pool of 11 spiking models using different random seeds. Each unique seed creates a distinct model instance, effectively representing an individual “subject” while keeping all other algorithmic hyperparameters constant. This approach allows us to determine if the model’s performance fluctuations mirror the diversity observed in human behavioral data. Each subject performed 4 trials of the Treasure Hunt Task. Each trial was randomly configured for: (1) the locations of the four tables within the arena, (2) the selection of 2 or 3 tables to contain objects, (3) the assignment of specific objects (selected from ten classes) to the non-empty tables, and (4) the traversal direction (clockwise or counter-clockwise) around the arena. In rare cases, our bipedal agent fell prior to visiting all (or any) chests. In these instances, any items the agent failed to encounter were excluded from the analysis. Consequently, our final results are based on 158 item-reveal events recorded across 41 successful trial segments.

The model’s behavior (Fig. 2) reproduced a key finding from the original experiment: accuracy increased with confidence (Fig. 2b; *ρ* = 0.48, *P* = 3.5 ×10^−7^, Spearman rank test, *n* = 96 queries). Several qualitative similarities with human behavior also emerged. Variability in mean accuracy across subjects mirrored human data (Fig. 2a,d), and as with humans, model accuracy distributions became increasingly concentrated near perfect scores with rising confidence, particularly around 0.9 (Fig. 2c,f). Nonetheless, some differences were apparent. The model was more accurate overall (mean = 0.85 ± 0.036) than humans (mean = 0.69 ± 0.017). Moreover, accuracy gains across confidence levels were uneven: a sharp increase from low to medium confidence contrasted with a modest gain from medium to high. Finally, while humans showed a slight bias toward higher scores even at low confidence, the model’s low-confidence outputs were skewed toward poor performance (accuracy ≤ 0.3). Together, these findings indicate an imperfect yet compelling correspondence between model and human behavior.

The neural data associated with the Treasure Hunt task focused on hippocampal local field potential (LFP) recordings during item presentation events. Differences in spectral power during these intervals were linked to outcomes in the later querying phase: with an increase in power in the left, but not right, hippocampal low-theta (1-3 Hz) band for items recalled over those not recalled being a key finding from the original study. In comparing to the LFP data, we targeted the Associative Memory (AM) subcomponent of the SSP-SLAM module, as it directly contributes to spatial learning in our model. We derived simulated LFPs by first convolving spike trains with a biphasic kernel to emulate transmembrane current fluctuations. Neurons were then spatially embedded relative to a hypothetical recording site, with distances sampled proportional to *r*^2^, as expected under uniform neuronal density. Extracellular potential was then calculated using the point-source model (Destexhe & Bedard, 2013) with *σ* = 1.7 µS m^−1^ assumed for extracellular conductivity. We replicate the spectral analyses described in the original study, and adopt the definition of recall as above-subject median accuracy and medium or high confidence, as in the original study.

Fig. 3 shows empirical (top row) and simulated (bottom row) of LFP time courses of spectral power across low theta (1–3 Hz), theta (3–10 Hz), and gamma (40–100 Hz) frequency bands, aligned to stimulus onset and split by recall outcome. A prominent feature among both simulated and empirical transients was a sharp increase in power at item onset. The model qualitatively reproduces the “left hippocampal memory effect” described by Miller et al., with higher simulated LFPs during the peri-stimulus window for recalled vs not recalled items. However, replicating the statistical analyses described in the original study revealed differences between the model and data. Notably, whereas in the empirical data a significant effect was confined to low-theta specifically, we observed memory effects across all bands (low theta: 1.26 ±0.38, theta: 1.28± 0.44, gamma: 1.91 ±1.15, all *P* < 0.001, one-sided, one-sample t-test).

**Figure 3:**
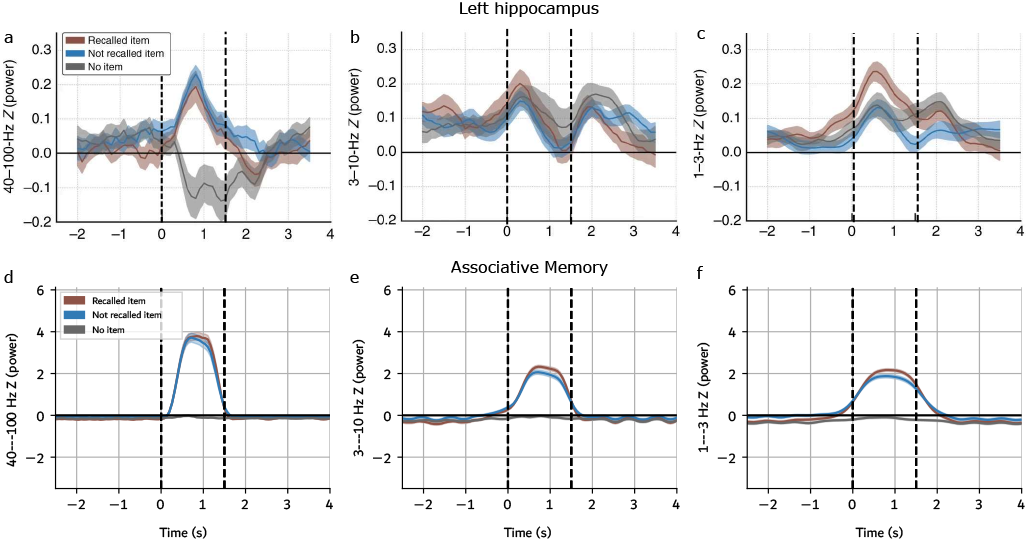
Spectral power time courses during item encoding from (top) human intracranial LFP recordings in the left hippocampus and (bottom) spike-derived LFP signals from the associative memory module. Chests open at *t* = 0 s to reveal an item (or none; gray) and disappear at *t* = 1.5 s. Recall is defined in the main text. Shaded regions indicate ±s.e.m. within each event category. Top row adapted from Miller et al. Fig. 5, under CC BY 4.0. Changes: Cropped, annotated.

Empirical hippocampal LFPs in the no-item condition showed qualitatively similar transients across all bands except gamma, where opening an empty chest produced a sharp power decrease. By contrast, model-derived LFPs for this control condition were largely flat, consistent with the model’s inability to encode any perceptual event at that location. Empirical LFPs also often exhibited a second post-stimulus transient absent from the model.

Collectively, these findings suggest that the dynamics within the left hippocampus during item presentation may facilitate the acquisition of place-object associations. While the model does not capture details of the empirical data—such as the post-stimulus transients—it demonstrates how a biologically plausible local learning rule can govern the formation of object identity representations within a spatial map, and captures the main effect in the human experiment, of LFPs distinguishing recalled vs. non-recalled items, with spiking neural mechanisms.

## Discussion

This work demonstrates the construction of an embodied map learning agent in a spiking neural network able to perform the ethologically-relevant Treasure Hunt task. Our model successfully acquires object-place associations from visual and velocity signals during learning, and leverages HRR operations to guide behavior in a subsequent querying phase-transforming between allocentric and egocentric frames for motoric pointing and computing subjective confidence along a three-point scale. Although raw human data were unavailable for distribution-level comparison, the model reproduced the main accuracy–confidence correlation, supporting a probabilistic interpretation of subjective confidence. Its spiking SLAM subsystem also produced dynamics resembling the left-hippocampal memory-related LFP effect.

Despite these advances, the model remains incomplete as a neuroethological account. Critically, symbolic queries bypass object recognition and short-term retention, demands which a truly embodied agent must meet. The modeled task omits visually guided navigation, distractors, trial-wise learning, and phase-dependent context switching, and most components remain rate-coded rather than spiking. Behavioral and neural discrepancies identify the main targets for refinement. The behavioral comparison identified a significant model/human accuracy gap. Systematic perturbations along the signal chain could identify constraints sufficient to bring performance into the human range, although parameter fitting would weaken the dataset as an independent benchmark. A critical unmodeled source of error is velocity representation: our agent receives perfect IsaacSim velocity input, whereas velocity-input noise is a key contributing factor to human path integration errors (Stangl et al., 2020). Neurally, the broadband memory effect may reflect higher model firing rates than observed *in vivo*, a possibility currently under quantitative sensitivity analysis. The absence of post-stimulus transients further suggests unmodeled neuronal or circuit mechanisms; future work should test whether adaptive neuron models or additional spiking subnetworks within SSP-SLAM can reproduce these effects. Comparisons of refined models should be based on a larger sample size, ideally on par with Miller et al. (2018).

Overall, the findings support biologically grounded embodied models for linking neural dynamics, internal belief, and overt behavior. As virtual-reality and mobile-recording methods mature (Stangl et al., 2023), such models will be increasingly important for capturing biological cognition.

## Acknowledgments

The authors would like to thank the anonymous reviewers for insightful comments that helped improve this paper. This work was supported by CFI (52479-10006) and OIT (35768) infrastructure funding as well as the Canada Research Chairs program, NSERC Discovery grant 261453, AFOSR grant FA9550-17-1-0644. We used generative AI tools (ChatGPT/GPT-based assistants by OpenAI and Claude by Anthropic) for limited writing support (polishing/summarizing), brainstorming, and code assistance (example snippets and debugging). All AI outputs were critically reviewed and edited as needed by the authors, who take full responsibility for the final work.

## References

Aharon, G., Sadot, M., & Yovel, Y. (2017). Bats use path integration rather than acoustic flow to assess flight distance along flyways. Current Biology, 27(23), 3650–3657.

Bekolay, T., Kolbeck, C., & Eliasmith, C. (2013). Simultaneous unsupervised and supervised learning of cognitive functions in biologically plausible spiking neural networks. Proceedings of the annual meeting of the cognitive science society, 35(35).

Benhamou, S. (1997). Path integration by swimming rats. Animal Behaviour, 54(2), 321–327. 10.1006/anbe.1996.0464

Destexhe, A., & Bedard, C. (2013). Local field potential. Scholarpedia, 8(8), 10713.

DeWolf, T., et al. (2020). ABR Control: A python library for robotic arm path planning and control. https://github.com/abr/abr_control

Doeller, C. F., Barry, C., & Burgess, N. (2010). Evidence for grid cells in a human memory network. Nature, 463(7281), 657–661.

Dumont, N. S.-Y., & Eliasmith, C. (2020). Accurate representation for spatial cognition using grid cells. CogSci.

Dumont, N. S.-Y., Furlong, P. M., Orchard, J., & Eliasmith, C. (2023). Exploiting semantic information in a spiking neural slam system. Frontiers in Neuroscience, 17, 1190515.

Durrant-Whyte, H., & Bailey, T. (2006). Simultaneous localization and mapping: Part i. IEEE robotics & automation magazine, 13(2), 99–110.

Dyer, F. C. (1991). Bees acquire route-based memories but not cognitive maps in a familiar landscape. Animal Behaviour, 41(2), 239–246.

Eliasmith, C. (2013). How to build a brain: A neural architecture for biological cognition. OUP USA.

Furlong, M., & Eliasmith, C. (2022). Fractional binding in vector symbolic architectures as quasi-probability statements. Proceedings of the annual meeting of the cognitive science society, 44(44).

Glad, I. K., Hjort, N. L., & Ushakov, N. G. (2003). Correction of density estimators that are not densities. Scandinavian Journal of Statistics, 30(2), 415–427.

Habashy, K. (2026). Associative memory for hyperdimensional spatial representations with geodesic flow matching.

Hafting, T., Fyhn, M., Molden, S., Moser, M.-B., & Moser, E. I. (2005). Microstructure of a spatial map in the entorhinal cortex. Nature, 436(7052), 801–806.

Jacobs, J., Weidemann, C. T., Miller, J. F., Solway, A., Burke, J. F., Wei, X.-X., Suthana, N., Sperling, M. R., Sharan, A. D., Fried, I., et al. (2013). Direct recordings of gridlike neuronal activity in human spatial navigation. Nature neuroscience, 16(9), 1188–1190.

Jocher, G. (2020, May). YOLOv5 by Ultralytics (Version 7.0). 10.5281/zenodo.3908559

Komer, B. (2020). Biologically inspired spatial representation [Doctoral dissertation, University of Waterloo].

Komer, B., Stewart, T. C., Voelker, A. R., & Eliasmith, C. (2019). A neural representation of continuous space using fractional binding. Proceedings of the Annual Meeting of the Cognitive Science Society, 41.

Lajoie, P.-Y., Ramtoula, B., Wu, F., & Beltrame, G. (2022). Towards collaborative simultaneous localization and mapping: A survey of the current research landscape. Field Robotics, 2, 971–1000.

Milford, M. J., Wyeth, G. F., & Prasser, D. (2004). Ratslam: A hippocampal model for simultaneous localization and mapping. IEEE International Conference on Robotics and Automation, 2004. Proceedings. ICRA’04. 2004, 1, 403–408.

Miller, J., Watrous, A. J., Tsitsiklis, M., Lee, S. A., Sheth, A., Schevon, C. A., Smith, E. H., Sperling, M. R., Sharan, A., Asadi-Pooya, A. A., et al. (2018). Lateralized hippocampal oscillations underlie distinct aspects of human spatial memory and navigation. Nature communications, 9(1), 2423.

Mittelstaedt, M. L., & Mittelstaedt, H. (2001). Idiothetic navigation in humans: Estimation of path length. Exp Brain Res, 139(3), 318–332. 10.1007/s002210100735

Pizzino, C. A. P., Costa, R. R., Mitchell, D., & Vargas, P. A. (2024). Neoslam: Long-term slam using computational models of the brain. Sensors, 24(4), 1143.

Plate, T. A. (1992). Holographic recurrent networks. In S. Hanson, J. Cowan, & C. Giles (Eds.), Advances in neural information processing systems (Vol. 5). Morgan-Kaufmann.

Safron, A., Çatal, O., & Verbelen, T. (2022). Generalized simultaneous localization and mapping (g-slam) as unification framework for natural and artificial intelligences: Towards reverse engineering the hippocampal/entorhinal system and principles of high-level cognition. Frontiers in Systems Neuroscience, 16, 787659.

Seeley, T. D., & Buhrman, S. C. (1999). Group decision making in swarms of honey bees. Behavioral Ecology and Sociobiology, 45(1), 19–31.

Shah, D., Osiński, B., Levine, S., et al. (2023). Lm-nav: Robotic navigation with large pre-trained models of language, vision, and action. Conference on robot learning, 492–504.

Silveira, L., Guth, F., Drews-Jr, P., Ballester, P., Machado, M., Codevilla, F., Duarte-Filho, N., & Botelho, S. (2015). An open-source bio-inspired solution to underwater slam. IFAC-PapersOnLine, 48(2), 212–217.

Stangl, M., Kanitscheider, I., Riemer, M., Fiete, I., & Wolbers, T. (2020). Sources of path integration error in young and aging humans. Nature communications, 11(1), 2626.

Stangl, M., Maoz, S. L., & Suthana, N. (2023). Mobile cognition: Imaging the human brain in the real world. Nature Reviews Neuroscience, 24(6), 347–362.

Steckel, J., & Peremans, H. (2013). Batslam: Simultaneous localization and mapping using biomimetic sonar. PloS one, 8(1), e54076.

Taheri, H., & Xia, Z. C. (2021). Slam; definition and evolution. Engineering Applications of Artificial Intelligence, 97, 104032.

Tolman, E. C. (1948). Cognitive maps in rats and men. Psychological Review, 55(4), 189–208. 10.1037/h0061626

Voelker, A. R. (2020). A short letter on the dot product between rotated fourier transforms. arXiv preprint arXiv:2007.13462.

Voelker, A. R., Crawford, E., & Eliasmith, C. (2014). Learning large-scale heteroassociative memories in spiking neurons. Unconventional computation and natural computation, 7(2014).

Wehner, R., Bleuler, S., Nievergelt, C., & Shah, D. (1990). Bees navigate by using vectors and routes rather than maps. Naturwissenschaften, 77(10), 479–482.

Wehner, R., Boyer, M., Loertscher, F., Sommer, S., & Menzi, U. (2006). Ant navigation: One-way routes rather than maps. Current biology, 16(1), 75–79.

Yartsev, M. M., Witter, M. P., & Ulanovsky, N. (2011). Grid cells without theta oscillations in the entorhinal cortex of bats. Nature, 479(7371), 103–107.

Zhu, S. L., Lakshminarasimhan, K. J., & Angelaki, D. E. (2023). Computational cross-species views of the hippocampal formation. Hippocampus, 33(5), 586–599.

